# Retinotopy constrains the topology of neural manifolds in macaque visual cortex

**DOI:** 10.64898/2025.12.22.695909

**Authors:** Ramin Ardalani, Anno C. Kurth, Aitor Morales-Gregorio

## Abstract

Cortical population activity often evolves along low-dimensional manifolds, shaped by external activity, task-demands and the underlying neuronal network structure. The structure of population activity has been studied across many brain areas and species. However, the influence of retinotopic organization in the structure of population activity of the primate visual cortex *in vivo* remains understudied. Here, we show that retinotopy in primary visual cortex (V1) determines the topological structure in neural population activity in the macaque. We analysed large-scale extracellular recordings from two macaques implanted with 896-channel Utah arrays in V1 performing a receptive field mapping task, with high-contrast moving bars. Isometric Mapping (isomap) applied to the neural population activity revealed a loop-like manifold. Persistent homology on 10-dimensional embeddings confirmed a single dominant one-dimensional loop. To test whether retinotopy alone can account for this structure, we constructed a minimal neural response model and simulated activity with the same stimuli as the macaque neural data. The model reproduced the loop-like topology observed in the brain. Taken together, the experimental and simulated data show that retinotopic maps naturally induce loop-like neural manifolds in response to moving stimuli, directly linking travelling waves and rotational dynamics. Our results provide a basis for further analysis on how neuronal populations encode and process visual information at the population level.

## Introduction

In many mammalian species, early visual processing is organized in a highly structured, retinotopic fashion^1,2^. Adjacent points in the visual scene evoke activity of neighbouring neurons across the cortical surface, providing a fundamental scaffold for visual information processing^3,4^. Retinotopy is explicit in the primary visual cortex (V1) and maintained across early extrastriate cortex^3^, whereas at higher ventral stages the mapping becomes less purely retinal, with V4 showing interleaved feature domains^5,6^ and IT exhibiting feature-based topographic organization^7,8^.

Neural activity evolving over time can be viewed as a trajectory in a high-dimensional state space, with each neuron representing an orthogonal axis in this space^9,10^. In practice, neural systems do not explore the full space, but instead are often constrained to some subset with a lower effective dimension: these subsets are often called neural manifolds^11–15^. Such manifolds have been shown to organize task and behavioural variables in several brain areas, such as context-dependent decisions in macaque prefrontal cortex^16^, hand movements in macaque motor cortex^12,13,17^, odour-space representations in mouse piriform cortex^18^, head-direction coding in mouse anterodorsal thalamic nucleus^15^, and spatial position in mouse hippocampus^19^. In vision, manifold structure has been characterized in the mouse visual cortex^20,21^ and, less extensively, in macaques^22,23^.

Because these manifolds can exhibit a complex global organization, their structure is naturally studied with tools from computational topology^15,22,24^. Persistent homology extracts qualitative features, such as connected components (zero-dimensional) and loops (one-dimensional), which can reveal the organizing principles of neural dynamics^25–27^. These tools provide a framework for testing how the topology of the stimulus-driven neural manifolds in early visual cortex are constrained by retinotopy^25,26^.

Despite detailed knowledge of the retinotopy in V1, how early spatial maps shape the topological features of neural population responses has not been explicitly studied for primate *in vivo* data. Most topological data analyses characterize intrinsic manifold structure without explicitly linking it to retinotopic or feature maps^26,28^. Recent population-level work in visual cortex has characterized neural response manifolds with rich geometry and topology, but their relation to the underlying retinotopic layout is typically only implicit^26,29^. More broadly, recent theoretical work has established a direct link between neural population activity and the underlying connectivity structure between neurons^30,31^. That retinotopy fundamentally constrains the topology and geometry of neural representations is intuitive from a theoretical point of view, especially when assuming a somewhat linear mapping from retina to cortex. However, neuronal dynamics are non-linear and direct *in vivo* evidence for this link has not been previously published.

Here, we investigate whether and how the retinotopic layout of early visual cortex imposes constraints on the geometry and topology of neural population responses. To address this question, we analyse spiking data from macaque V1^32^. Using the non-linear dimensionality reduction method isometric mapping (isomap)^33^ we find a stable structure in the population activity in the form of two quasi-orthogonal loops. Using topological data analysis, we confirm that the double-loop topology of the neural response manifolds is not a dimensionality reduction artifact. We reproduce the experimental observations in a minimal visual response model with retinotopic input, directly linking the observed neural manifolds to retinotopy. Taken together, our results show that retinotopic maps in early visual cortex constrain the topological structure of the visual cortex population activity in primates.

## Results

### Neural responses to moving bars in the macaque visual cortex

To study the effect of retinotopy on population activity in the visual cortex, we analysed data from two male macaques (*N* = 2) implanted with 14 Utah arrays in V1, providing simultaneous recordings from 896 channels^32^. In the experiment, the animals fixated a central point while a high-contrast light bar appeared at a peripheral location and swept across the screen for 1000 ms in one of the four cardinal directions (rightward, leftward, upward, downward; illustrated in Figure 1A,B). On each trial, the animals were required to maintain fixation until bar offset to receive a reward, yielding well-controlled epochs of visually driven activity (Figure 1C). The raw extracellular electrophysiology signals from V1 were processed to detect single-unit activity (SUA); automated spike sorting and quality control identified 513 SUA in macaque L (Figure 1E) and 186 SUA in macaque A (Figure S1E). See Electrophysiological data from macaques L & A and Spike sorting for detailed methods on data acquisition and processing.

**Figure 1:**
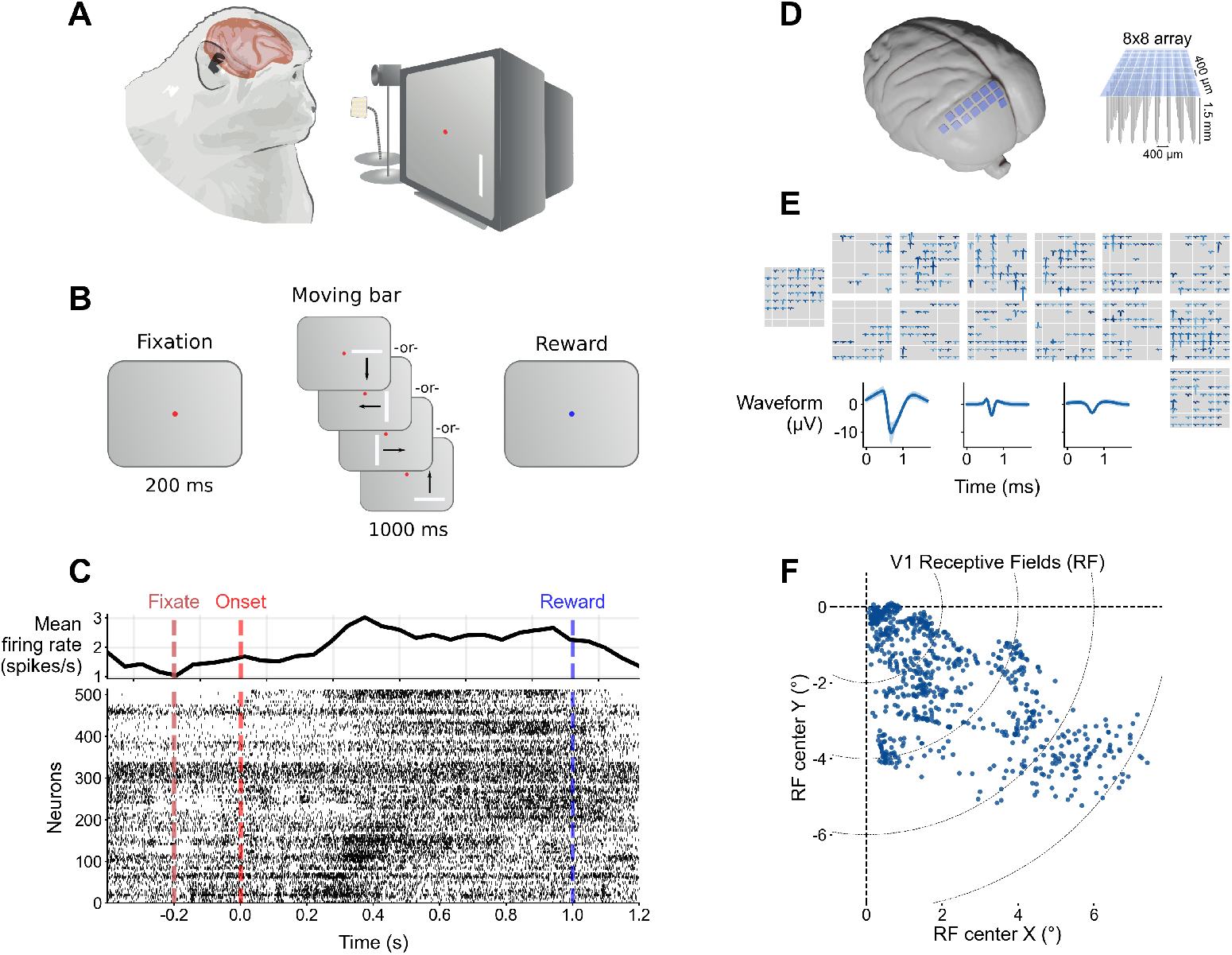
Neural responses to moving bars in the macaque visual cortex; data from Macaque L shown. **A)** Diagram of macaque and screen setup. **B)** Diagram of moving bar trials. **C)** Spike time raster plot for all V1 neurons for one example trial (bar moving downward), key times marked with vertical lines. Average firing rate across all neurons shown in upper sub-panel. **D)** Schematic of implant locations; 14 Utah arrays were implanted in V1 on the left hemisphere. **E)** Average spike waveforms for all the detected neurons in their approximate position. Bottom insets show the mean (thick line) and standard deviation (shading) of three example units. **F)** Receptive field location of all V1 electrodes covering the lower right quadrant of the visual scene.

The V1 arrays provided dense coverage of the lower-right quadrant of the visual scene (Figure 1F), such that the moving bar sequentially traversed the receptive fields of many simultaneously recorded SUA (Figure 1F). This is illustrated in an example trial with a downward-moving bar, where neuronal spike rates visibly increase at different times as the stimulus crosses their receptive fields (Figure 1C). These recordings yield a large-scale dataset of direction-labelled population responses, which form the basis for our subsequent analyses.

### Population responses in macaque V1 have a loop-like topology

To characterize the structure of V1 population activity, we analysed the geometry and topology of stimulus-evoked neural trajectories in state space. Because the moving bar is parametrized by a low-dimensional input (its position in a two-dimensional visual field, evolving smoothly over time), we expect the corresponding population responses to lie on a low-dimensional manifold. We therefore embedded the high-dimensional population activity into a shared three-dimensional space using Isometric Mapping (isomap). Isomap constructs a nearest-neighbour graph and seeks an embedding that approximately preserves geodesic distances along the data manifold, making it well suited for recovering curved low-dimensional structure from high-dimensional activity patterns. See Neural manifolds for details.

In this embedding, isomap revealed direction-specific trajectories that sweep along loop-like paths during stimulus presentation (Figure 2 for macaque L and Figure S1 for macaque A). During the pre-stimulus baseline, population activity clustered near a common “central” region of the state space. During each trial, activity departed from this central region, traced a loop-like excursion, and returned close to the centre at the end of the stimulus. Stimuli with the same bar orientation but opposite motion directions (e.g., leftward vs. rightward) produced highly similar loops traversed in opposite directions (Figure 2A,B and Figure S1A,B; time course indicated by coloured arrows). Changing the bar orientation (horizontal instead of vertical) yielded approximately orthogonal loops in the embedded state space (Figure 2C,D and Figure S1C,D). The consistency of these trajectories across repeated trials and stimulus conditions suggests a stable low-dimensional organization of V1 population activity, consistent within and across both studied macaques L and A.

**Figure 2:**
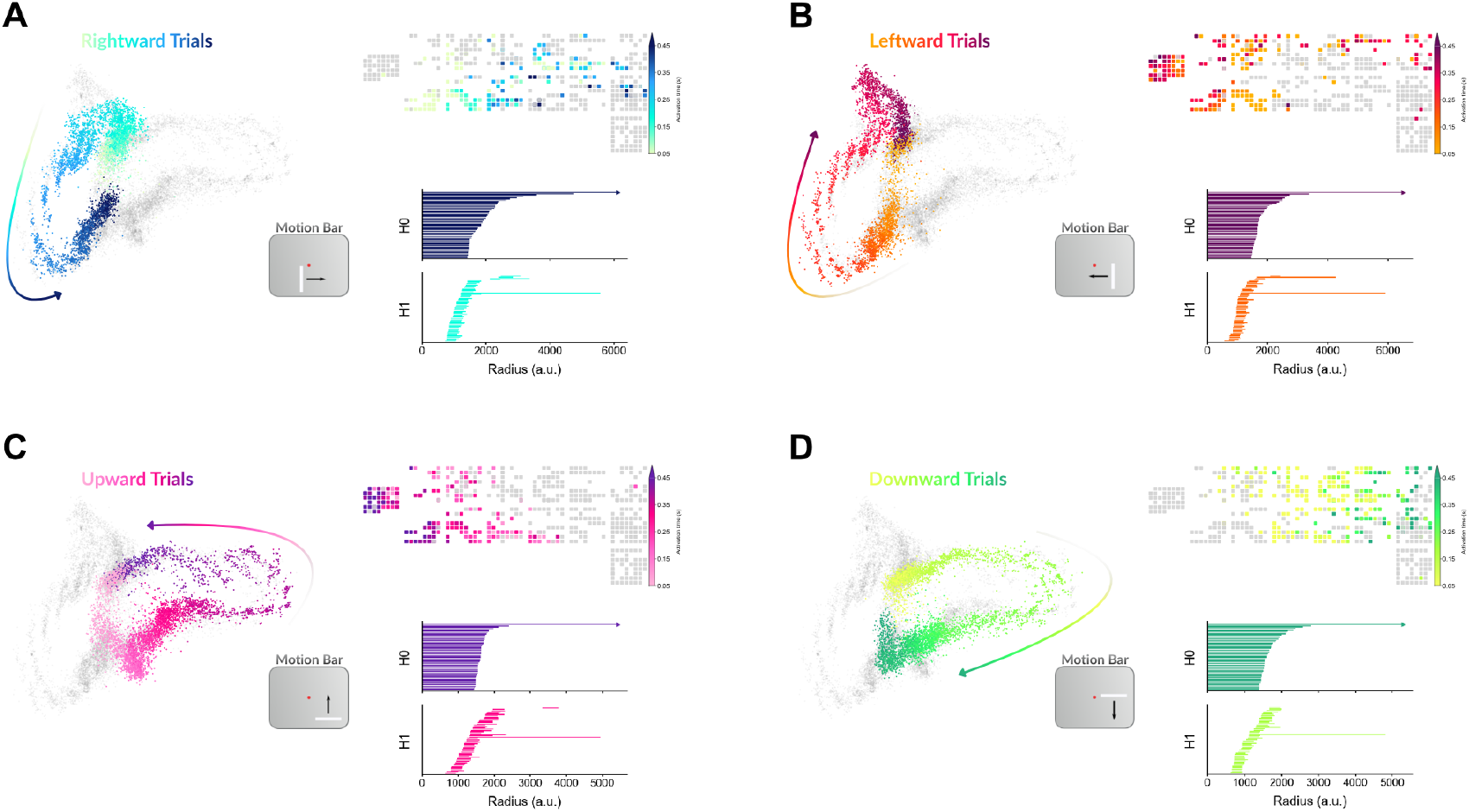
Neural activity in macaque V1 exhibits direction-dependent neural manifold and a loop-like topology. Results shown for **A)** rightward, **B)** leftward, **C)** upward and **D)** downward trials. Within each panel, (*Left*) 3-dimensional isomap embedding of the neural population activity discretized with 50 ms bins. The population responses unfold along a compact and loop-like trajectory. Colour indicates the time relative to trial onset, indicating smooth and consistent population dynamics across trials. (*Upper right*) Spatial location of the neurons and their activation latency with respect to trial onset. The activation-latency map highlights a sequential recruitment of recording sites that follows the expected retinotopic ordering. (*Lower right*) Vietoris–Rips persistence barcodes of the ten-dimensional isomap embedding of the neural population activity for the *H*_0_ and *H*_1_ homology groups. The barcodes indicate there is one *H*_0_ component, confirming there is a single continuous manifold, and one long-lived *H*_1_ component, confirming a high-dimensional loop topology.

To confirm that the loops observed in the isomap embedding are inherent to the high-dimensional data and not an artifact of dimensionality reduction, we use persistent homology. We apply persistent homology to each trial type separately. If the trials were analysed in combination, the large overlap between rightward–leftward or upward–downward trial manifolds would obfuscate the interpretation of the results. Persistence barcodes show rapid merging into a single connected component (*H*_0_), which indicates that the population activity is confined to a single fully connected component and is not composed of multiple disjoint manifolds. For all trial types, we observed a long-lived *H*_1_ barcode over a broad range of radii, indicating a loop-like topology with a one-dimensional hole (Figure 2A–D and Figure S1A–D lower right sub-panels). Thus, the persistence barcodes confirm the loop-like topology of the neural population activity in response to the moving bar stimuli.

### Simulated retinotopic responses reproduce neural topologies

We suggest that the simplest interpretation of the observed loop-like neural manifolds is retinotopy, due to the orderly and overlapping activation of adjacent units producing a loop in state space. To test this hypothesis, we simulated visual responses as a moving Gaussian activity wave with white noise over a grid of uniformly distributed neurons. Simulations were performed with stimuli moving in the four cardinal directions, replicating the experimental conditions in the macaques.

Schematic illustrations of the simulated responses are shown for each trial type in Figure 3. Note that our simulations directly model the neural responses, ignoring recurrent interactions between neurons and their non-linear dynamics. We find that the simulated responses reproduce the loop-like structure of population activity (Figure 3). On the one hand, the isomap embeddings of the simulated responses display nearly perfect overlap of the loops for trials with the same stimulus orientation but opposite moving directions. On the other hand, trials with orthogonal stimulus orientation display almost perfectly orthogonal loops. The existence of the loop-like topology in the simulated data is confirmed by persistent homology, with one long-lived *H*_1_ component present, analysed separately for each trial type.

**Figure 3:**
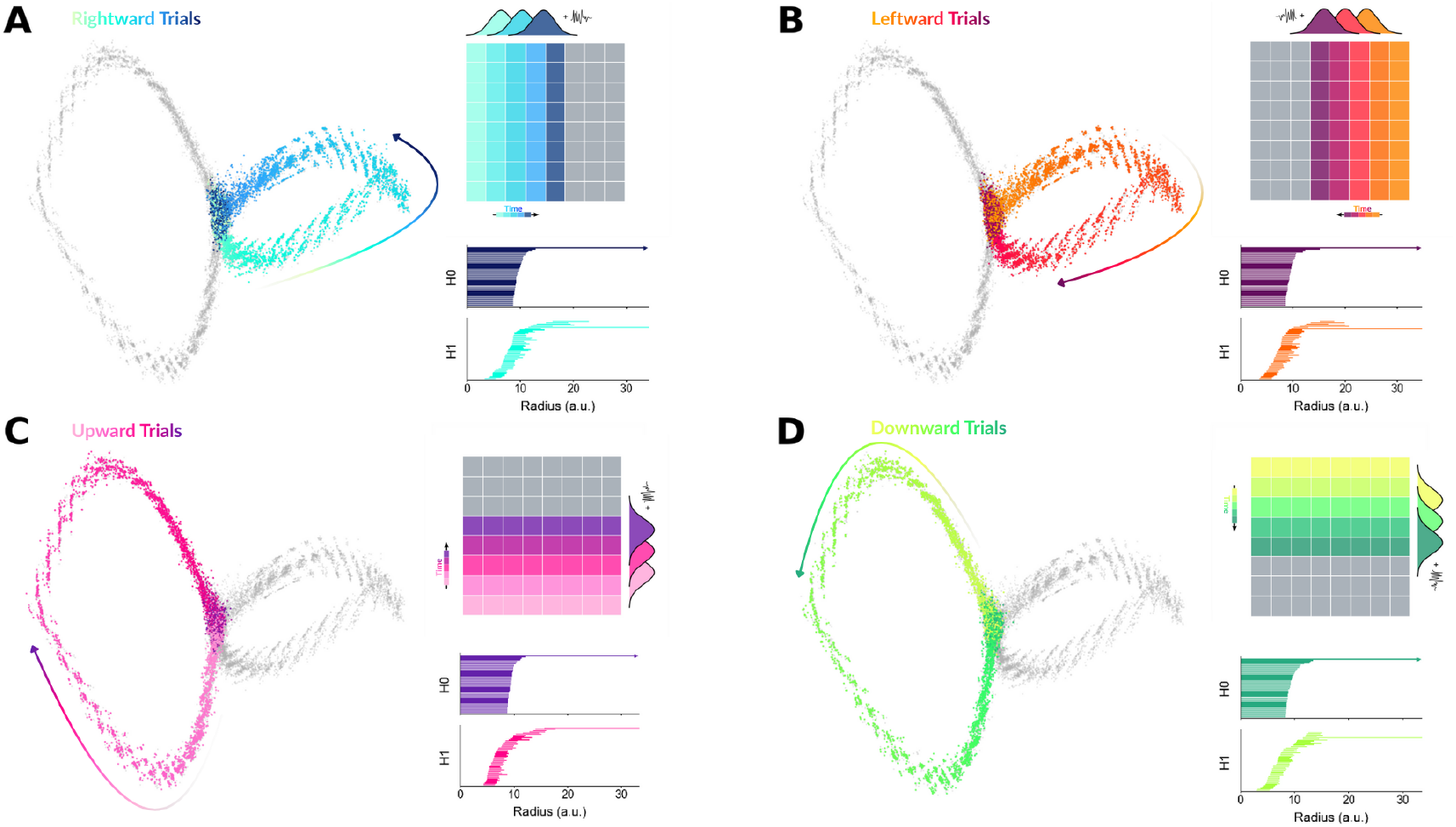
Simulated retinotopic activity reproduces direction-specific neural manifold topology. Results are shown for **A)** rightward, **B)** leftward, **C)** upward, and **D)** downward trials. Within each panel, (*Left*) simulated population activity embedded in a low-dimensional space using isomap, coloured by within-trial time. The trajectory forms a smooth loop-like path, indicating consistent temporal evolution along a confined manifold. (*Upper right*) the activation-latency map across the simulated recording grid, demonstrating an orderly spatio-temporal progression due to the retinotopic layout of the model. (*Lower right*) Vietoris–Rips persistence barcodes of the ten-dimensional isomap embedding of the simulated population activity. The barcodes indicate that there is one *H*_0_ component, confirming a single connected manifold, and one long-lived *H*_1_ component, confirming a high-dimensional loop topology.

All in all, the simulated data confirm that the observed topologies in macaque V1 can emerge from retinotopy.

## Discussion

Our analyses show that population activity in V1 during a simple moving-bar stimulus evolves along loop-like manifolds. For each motion direction, the population trajectory forms a one-dimensional loop in state space, both in macaque V1 recordings (Figure 2) and in a minimal retinotopic simulation (Figure 3). Taken together, our results suggest that the loop topology arises from the sequential recruitment of retinotopically organized neurons as the moving bar traverses the visual scene. This generates a stimulus-driven spatio-temporal travelling wave across cortex^34,35^, which follow loop-like topology in the neuronal state space^36^.

Our results connect to the broader literature on rotational dynamics and low-dimensional neural manifolds. In motor cortex, rotational trajectories during reaching and related behaviours are observed, motivating a dynamical-systems interpretations of movement generation^17,37–39^. Furthermore, rotational structure is ubiquitous across brain areas and different tasks^10^, suggesting that it can arise from generic properties of population responses. In particular, sequential activation of neurons can generate apparent rotations in state space^36,40^. Furthermore, neurons arranged in a ring networks also exhibit rotational dynamics in state space^41^. Thus, rotational trajectories in state space are the direct consequence of spatially organized travelling waves, but can also arise from non-spatially organized repeated sequential activation of neurons. Our V1 results fit naturally within this perspective: the loop trajectories observed here are well explained by an orderly stimulus-driven progression across a retinotopic map, rather than requiring some intrinsic oscillatory generator or other recurrent network effects.

A key practical implication is that the observed topology depends on how broadly the recorded population tiles visual space. In our dataset, the moving bar sweeps across a large portion of the visual field representation because the 4 *×* 4 *mm* Utah arrays span a wide range of receptive-field locations within V1 (Figure 1). Under this broad sampling, the stimulus-evoked activity progresses smoothly across the array and the population response closes into a loop (Figure 2). By contrast, if one recorded from a much narrower cluster of receptive fields, the same stimulus would be expected to capture only a short segment of the travelling wave, producing an short and open arc rather than a smooth closed loop. This provides a concrete, sampling-dependent prediction: restricting recordings to a spatially compact subregion could weaken or eliminate the loop’s topological signature even if the underlying cortical activation remains wave-like.

The simulation results further support the retinotopic origin of the experimental observations (Figure 3). Without specifying an explicit dynamical system or recurrent network interactions, a feedforward retinotopic model reproduces the main qualitative features of the experimental data: direction-dependent loop trajectories, overlapping trajectories for opposite stimulus movement direction, and approximate orthogonality between loops elicited by orthogonal stimulus orientations. This agreement indicates that retinotopy is sufficient to generate the observed loop topology for the moving-bar stimulus. At the same time, this does not rule out additional contributions from recurrent circuitry, gain control, or contextual modulation in real V1. Instead, the simulations suggest there is a retinotopy-driven backbone which can be further modulated by additional mechanisms.

Retinotopy is not the only organizing principle in V1. Across many mammalian species, especially primates and humans, neurons are arranged into maps and columns selective for orientation, motion direction, ocular dominance, and colour^42–45^. These feature maps are largely embedded within the retinotopic layout and can strongly shape local response patterns, influencing neuronal variability and correlations^46,47^. Our results with high-contrast moving bars suggest that retinotopy provides the scaffold for the topology of neural manifolds, while the selectivity for other features is expected to modulate the loop’s detailed geometry (e.g., thickness, curvature, and orientation in embedding space) without necessarily changing its one-dimensional topology. For example, direction- and orientation-tuned neurons may amplify different parts of the trajectory depending on motion direction, producing systematic deformations while preserving the overall loop.

Naturalistic vision is far richer than simple moving bars, leading to higher-dimensional neuronal responses than simple bars or gratings^29^. Studies of population activity at a single-neuron resolution in response to naturalistic images mostly focus in the mouse visual cortex^20,48,49^, with few studies focusing on primates^22,50^. Images appear to be encoded by neurons on a smooth continuous manifold in human neuroimaging recordings^48,51,52^. The geometry and dimensionality of those representations is progressively reshaped along the visual hierarchy, both in the brain and in deep neural networks trained for vision^53–55^. Our work suggests the topology of the neural representations is constrained by the retinotopy and other feature maps, which will likely also extend to higher visual areas due to the topographic connectivity along the hierarchy^56^.

In summary, the topology of V1 activity during moving-bar stimulation is constrained by the retinotopy: a smooth sweep of activity across V1 yields loop-like neural trajectories. This conclusion is supported by macaque recordings and by a minimal model, indicating that the observed loops do not reflect a finely tuned recurrent oscillator, and correspond to the neural representation of the visual stimulus. More generally, our results suggest a scale separation in how V1 structure appears in population activity: retinotopy provides a global scaffold determining the topology of stimulus-evoked trajectories, whereas feature selectivity likely modulates the local geometric detail.

## Methods

### Electrophysiological data from macaques L & A

We used publicly available neural activity data^32^ recorded from the neocortex of rhesus macaques (*N* = 2) during a visual task. The macaques were implanted with 14 8×8 Utah arrays (Blackrock Microsystems) in the primary visual cortex (V1) for a total of 896 electrodes. The electrodes were 1.5 mm long, thus recording from deeper layers of the gray matter, likely layer 5. The system recorded the electrical potential at each electrode with a sampling rate of 30 kHz. A full description of the experimental setup and data collection and preprocessing has already been published^32^; here we provide only the details relevant to this study.

The macaques were trained to fixate their gaze at the centre of a screen during the entire trial. A single thin white moving bar was presented on the screen, either horizontal or vertical. The bars moved in four possible directions: left-to-right or right-to-left for vertical bars, and up-to-down or down-to-up for horizontal bars. In experiments with both macaques, the moving bars were 20 degrees of visual angle (*dva*) in length, 0.19 *dva* in thickness, and moved at 20 *dva/s*. This task was performed to map the receptive fields (RFs) of the recorded neurons; the methods used are explained elsewhere^32^, and the estimated RFs are shown in Figure 1F.

### Spike sorting

The raw data for all sessions were spike-sorted using a semi-automatic workflow with custom preprocessing implemented in `spikeinterface`^57^ and `Kilosort4`^58^.

First, we band-pass filtered the 30 kHz raw signals between 0.5 kHz and 6 kHz using a fourth-order Butterworth filter. Second, the signals were whitened to remove spurious correlations originating from crosstalk, as described in Oberste-Frielinghaus et al.^59^. The preprocessed signals were then passed to the Python implementation of `Kilosort4`^58^ to perform spike sorting. Briefly, `Kilosort4` detects spike events using a template-matching approach, then clusters the templates, which are finally merged based on inter-spike intervals (ISI) and other metrics. We deactivated all preprocessing steps performed by `Kilosort4`, especially drift correction, which is designed for acute laminar probes and yielded undesirable results for the two-dimensional Utah arrays.

After automatic sorting, waveform clusters were evaluated using a range of quality metrics, including firing rate, waveform signal-to-noise ratio (wfSNR), ISI, presence ratio, and locking to the 50 Hz electric grid noise. The results were saved in the NIX format alongside all relevant metadata, such as electrode positions, cortical area, and array ID.

### Neural manifolds

Spike trains from macaque V1 were binned in 50 ms windows to form a time-by-neuron matrix. Note that the results were qualitatively equivalent both for shorter and longer time bins (not shown).

Some time points lie far from the bulk in state space, which we attribute to noise and therefore sought to remove. To identify outliers, we used a density-based rule similar to that of Chaudhuri et al.^15^. First, we computed the pairwise Euclidean distance matrix across time points and took the 10th-percentile value from the distance distribution *D*_10_. Second, we counted the number of neighbours for each point within distance *D*_10_ and discarded the bottom 10% of points based on this neighbour count. The isomap embedding was fit on the cleaned set and applied to the full time series.

We then applied isomap^33^ using five nearest neighbours for the geodesic distance calculation. We fit isomap to the cleaned data without the outliers, but then we projected all data points (including outliers) into a three-dimensional space for visualization and a ten-dimensional space for computing the topological features.

Trials were aligned to stimulus onset and grouped by motion direction (rightward, leftward, upward, downward). For each group, we extracted the corresponding segments from the embeddings and concatenated them into continuous trajectories across trials. These trajectories serve as the neural manifolds shown in Figure 2, Figure S1 and were used for the topological data analysis.

### Topological data analysis

We used persistent homology to verify that the lower-dimensional structure seen in the three-dimensional projections reflects genuine topology rather than an artifact of dimensionality reduction. Before computing persistence barcodes, we embedded the time-indexed population activity into a ten-dimensional subspace using isomap^33^, which approximately preserves geodesic distances on the manifold and is therefore suitable for topological analysis.

To compute the Vietoris–Rips persistence barcodes of the neural manifold, we used an efficient open-source implementation (https://ripser.scikit-tda.org). Briefly, the filtration inflates balls of radius *r* around each point and includes a (*k* − 1)-simplex whenever *k* points are pair-wise within distance *r*. This yields a simplicial complex whose topological features summarize the manifold. We computed barcodes up to homology dimension one, i.e., *H*_0_ (connected components) and *H*_1_ (one-dimensional loops), and for clarity we visualize only the 50 longest bars for each trial type and homology group. The analysis was performed separately for each motion direction on the ten-dimensional isomap trajectories.

### Simulation of neural responses

We simulated neural responses using a minimal feedforward model. We generated synthetic population activity from 100 simulated neurons arranged on a 10 *×* 10 square grid with coordinates *x, y* ∈ [−9, 9] a.u., mimicking a retinotopic layout. The moving bar responses in the visual cortex were modelled as a Gaussian bell with a time-varying mean *µ*(*t*), variance *σ*^2^ = 0.15 and amplitude 2, corresponding to neural activity sweeping across the 2-dimensional grid of synthetic neurons. The Gaussian bell was swept in the four cardinal directions (rightward, leftward, upward, downward), to reproduce the same experimental conditions as in the animal experiments. Note that the variance of the Gaussian had to be large enough to activate multiple adjacent neurons, otherwise the activity would not display the loop-like topology, instead going back-and-forth from the origin at each time point (not shown).

In all cases, uniform noise *η* ∈ [0, 0.15] was added to each synthetic neuron at every time step. The noise was not necessary to reproduce the loop-like topology, but was introduced to increase the geometric resemblance to real neuronal responses by increasing the “thickness” of the loops. Using white noise or other uncorrelated noise sources instead of uniform noise yielded no qualitative difference to the simulation results (not shown).

Importantly, this simulation does not model a recurrent network. The responses are purely stimulus-driven due to the retinotopy-like arrangement of the simulated neurons. We did not simulate the interactions between neurons in a network.

The simulated responses were then processes following the same dimensionality reduction and persistence homology methods applied to the experimental data. All spatial and temporal units were arbitrary for the simulated data.

## Data availability

The raw data used in this study are publicly available in a GIN repository under https://doi.org/10.12751/g-node.i20kyh, which was previously published^32^.

The spike-sorted data and additional files required to reproduce the results of this study will be made available upon publication in a peer-reviewed journal.

## Code availability

The Python scripts used for preprocessing, analysis, and plotting the results of this paper will be made available upon publication in a peer-reviewed journal.

## Author contributions

Conceptualization: AMG, AK; Data curation: AMG; Data preprocessing: AMG; Methodology: RA, AK, AMG; Software: RA; Formal analysis: RA; Visualization: RA, AMG; Writing—original draft: RA; Writing—review & editing: RA, AK, AMG; Supervision: AMG; Resources: AMG; Funding acquisition: AMG.

## Acknowledgments

We thank Jon Martinez Corral for providing feedback on our manuscript.

This work received funding from the Programme Johannes Amos Comenius (OP JAK) under the project “MSCA Fellowships CZ–UK3” (registration number CZ.02.01.01/00/22 010/0008220).

## Supplementary materials

**Figure S1:**
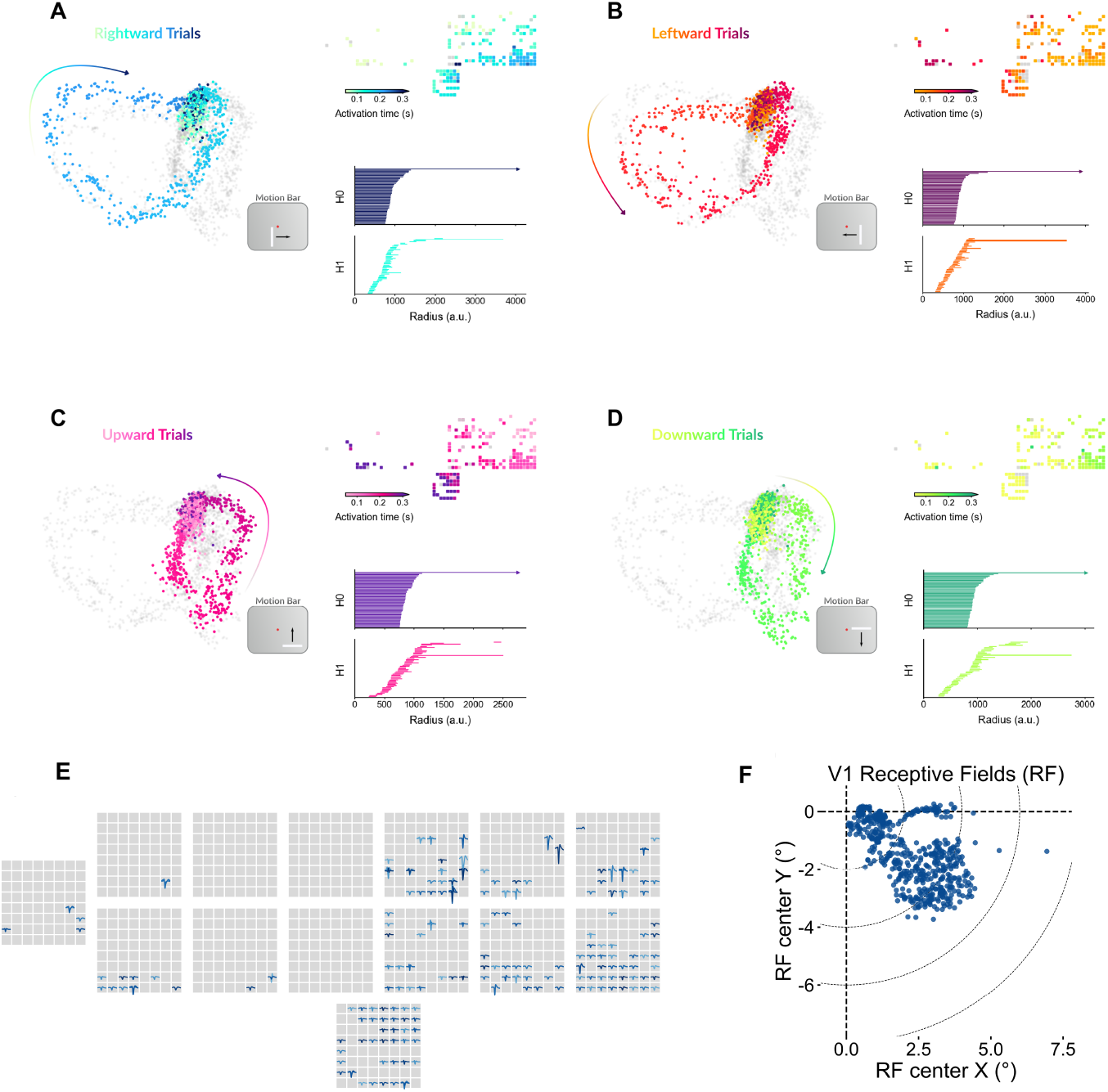
Neural activity in V1 exhibits direction-dependent neural manifolds and a loop-like topology in macaque. **A**. Results are shown for **A)** rightward, **B)** leftward, **C)** upward, and **D)** downward trials. Within each panel, (*Left*) three-dimensional isomap embedding of the neural population activity discretized into 50 ms bins. The population responses unfold along a compact and loop-like trajectory. colour indicates time relative to trial onset, indicating smooth and consistent population dynamics across trials. (*Upper right*) spatial locations of the neurons and their activation latency with respect to trial onset. The activation-latency map highlights a sequential recruitment of recording sites that follows the expected retinotopic ordering. (*Lower right*) Vietoris–Rips persistence barcodes of the ten-dimensional isomap embedding of the neural population activity for the *H*_0_ and *H*_1_ homology groups. These analyses demonstrate that the loop-like topology is present in two different subjects, confirming its generality. **E)** Average spike waveforms for all the detected neurons in their approximate position for macaque A. **F)** Receptive field location of all V1 electrodes covering the lower right quadrant of the visual scene.

